# Broadband Tuning the Voltage Dependence of a Sodium Channel

**DOI:** 10.1101/2020.11.21.392571

**Authors:** Eedann McCord, Goragot Wisedchaisri, William A. Catterall

**Affiliations:** Department of Pharmacology, University of Washington, Seattle, WA 98195-7280 USA

## Abstract

Voltage-gated sodium channels initiate action potentials in prokaryotes and in many eukaryotic cells, including vertebrate nerve and muscle. Their activation is steeply voltage-dependent, but it is unclear how the voltage sensitivity is set or whether it can be broadly shifted to positive voltages. Here we show that the voltage dependence of activation (V_A_) of the ancestral bacterial sodium channel Na_V_Ab can be progressively shifted from −118 mV to +35 mV in chimeras with increasing numbers of amino acid residues from the extracellular half of the voltage sensor of human Na_V_1.7 channels. In a minimal chimera in which only 32 residues were transferred, we analyzed the effects of six additional mutations of conserved amino acid residues singly, in pairs, and as triple mutations. The resulting chimeric mutants exhibited a broad range of voltage sensitivity from V_A_=−118 mV to V_A_=+120 mV. Three mutations (N48K, L112A, and M119V) shifted V_A_ to +61 mV when substituted in Na_V_Ab itself, and substitution of two additional Cys residues in the Cys-free background of Na_V_Ab further shifted V_A_ to +105 mV. In these mutants, measurement of gating currents revealed that the voltage dependence of gating charge movement (V_Q_) shifted to positive membrane potentials as much or more than V_A_, confirming that the gating charges are trapped in their resting positions by these V_A_-shifting mutations. Our results demonstrate broadband shifting of V_A_ and V_Q_ of a sodium channel across a range of 240 mV and provide a toolbox of methods and constructs to analyze sodium channel structure and function in the resting state at 0 mV and in activated states at positive membrane potentials.

**GRAPHICAL ABSTRACT:** The complete range of broadband tuning of voltage-dependent activation of a sodium channel.

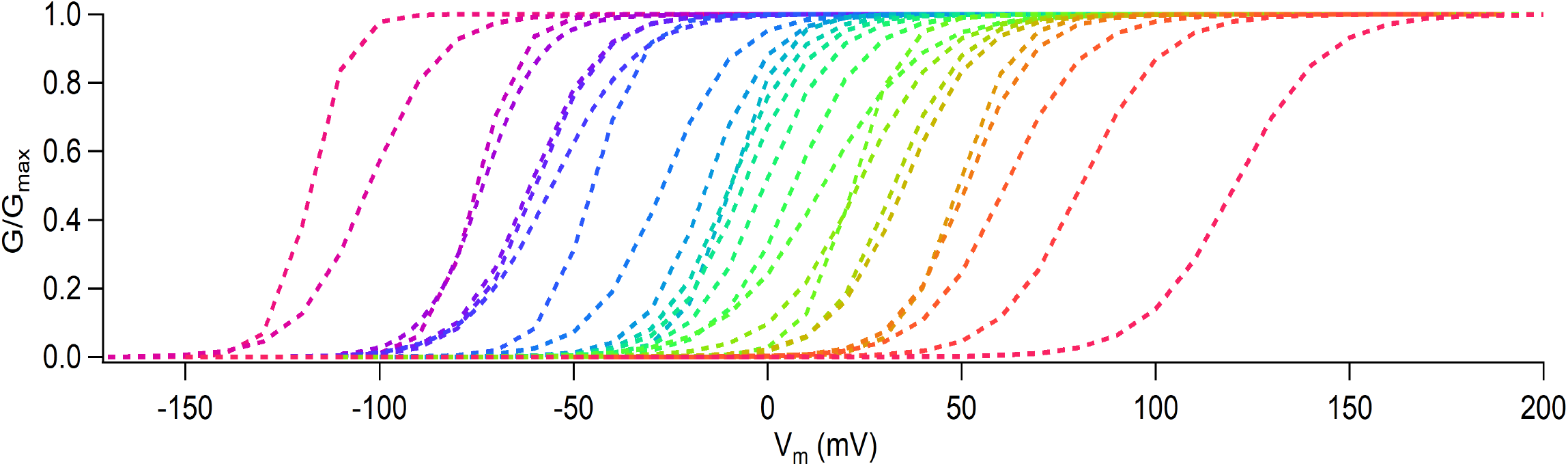

## INTRODUCTION

Animal life depends on electrical signaling via action potentials. The swift conduction of action potentials along the cell surface of neurons, myocytes, and other excitable cells confers the ability to coordinate rapid events – allowing animals to breathe, move, and think (Hille, 2001; Hodgkin and Huxley, 1952). Action potentials are initiated by voltage-gated sodium channels, which open in response to membrane depolarization and then inactivate within 1-5 msec (Hodgkin and Huxley, 1952). Depolarization of the cell membrane also activates voltage-gated potassium channels, which open more slowly and conduct potassium out of the cell to repolarize the membrane potential (Hodgkin and Huxley, 1952). In order to explain the voltage dependence of activation of sodium channels, Hodgkin and Huxley proposed that membrane-bound charges associated with sodium channels respond to the change in electric field, move across the membrane upon depolarization, and return to their resting position upon repolarization. Based on the steepness of voltage-dependent activation, they proposed that 3 singly charged ‘gating particles’ control activation and one gating particle controls inactivation (Hodgkin and Huxley, 1952). Their prescient prediction was confirmed when the movement of these gating particles was directly measured in the form of gating currents, which are nonlinear capacitive currents that arise from the movement of the gating particles (Armstrong and Bezanilla, 1973, 1977; Bezanilla and Armstrong, 1977). Fast, outward movement of positive gating charge was triggered by depolarization (ON gating currents), and a corresponding inward OFF gating current was observed upon hyperpolarization of the membrane. These gating currents are the electrical signature of the voltage-dependent activation process.

The protein components of sodium channels were initially identified by photolabeling studies using scorpion toxins, which revealed a complex of an α subunit and a smaller β subunit (Beneski and Catterall, 1980). Sodium channels were purified from electric eel electroplax (Barchi, 1983; Miller et al., 1983), skeletal muscle (Barchi, 1983), and brain (Hartshorne and Catterall, 1981, 1984; Hartshorne et al., 1982). The brain sodium channel was reconstituted into phospholipid bilayers and vesicles, and measurement of ion flux and single-channel currents verified that these purified proteins were in fact voltage-gated sodium channels (Hartshorne et al., 1985; Talvenheimo et al., 1982; Tamkun et al., 1984). Cloning and sequencing cDNA encoding sodium channel α subunits revealed a polypeptide of ~2000 amino acid residues having 24 transmembrane segments organized in six homologous domains (Noda et al., 1986; Noda et al., 1984). Extensive structure-function studies showed that the first four helices (S1-S4) of each subunit comprise the voltage-sensing domain (VS), while the fifth and sixth (S5-S6) transmembrane helices from all four subunits come together to form the pore module (PM) (Catterall, 2000). The S4 segment contains a repeated motif of a positively charged residue flanked on each side by hydrophobic ones in a three-residue pattern that is repeated 4-8 times in sodium channel VS. Mutagenesis studies revealed that these charged residues on the S4 segment serve as the gating charges that enable the VS to sense fluctuations in membrane potential and respond by opening the pore (Bezanilla, 2000; Catterall, 2000; Kontis et al., 1997; Stuhmer et al., 1989). Chemical labeling, voltage-clamp fluorometry, and disulfide-crosslinking studies revealed that changes in membrane potential drive the S4 segments of sodium channels across the membrane through a narrow hydrophobic seal, catalyzed by interactions with negative charges on the intracellular and extracellular sides (Chanda and Bezanilla, 2002; DeCaen et al., 2011; DeCaen et al., 2009; DeCaen et al., 2008; Yang et al., 1996; Yang and Horn, 1995; Yarov-Yarovoy et al., 2012). A “sliding-helix” model in which the S4 segment moves outward through the VS upon depolarization and exchanges ion pair partners has emerged as a consensus mechanism for voltage sensing (Catterall, 1986; Catterall et al., 2017; Vargas et al., 2012; Wisedchaisri et al., 2019; Yarov-Yarovoy et al., 2012).

The unexpected discovery of bacterial sodium channels showed that they are homotetrameric complexes of four identical subunits with six transmembrane segments (Ren et al., 2001). This simple organization facilitated structural studies using X-ray crystallography, which defined the structure of the bacterial sodium channel Na_V_Ab at 2.7 Å resolution (Payandeh et al., 2011). Other bacterial sodium channels have similar structures (Ahern et al., 2016; Catterall et al., 2017; McCusker et al., 2012; Zhang et al., 2012). Moreover, eukaryotic sodium channels analyzed by cryo-electron microscopy (cryo-EM) have a similar core structure, with RMSD in the range of 3 Å for the transmembrane regions of mammalian sodium channels from skeletal muscle, nerve, and heart (Jiang et al., 2020; Pan et al., 2019; Pan et al., 2018; Shen et al., 2019). These structural results enabled analysis of the molecular basis of voltage-dependent gating at the atomic level, but the range of functional states observed in these structural studies was limited because they were all carried out in the absence of a membrane potential where partially activated and steady-state inactivated states of sodium channels are most stable. Here we used a combination of structure-guided mutagenesis and biophysical analysis to tune the voltage dependence of sodium channels over a broad range and determine whether they can function normally in the positive membrane potential range. Our results reveal that broadband tuning of the voltage dependence of a sodium channel over a range up to 240 mV is possible (see Graphical Abstract), and they provide a novel approach to analysis of the structure and function of sodium channels in multiple states that normally would not be accessible for study by structural biology methods at 0 mV. As proof of this concept, we have used one of the constructs described here to determine the structure of the resting state of a sodium channel at high resolution (Wisedchaisri et al., 2019).

## MATERIALS AND METHODS

### Construction of Na_V_Ab/Na_V_1.7-VS2 chimera and mutants

Na_V_Ab/Na_V_1.7-VS2 cDNAs with a wide range of voltage-shifting mutations and combinations were constructed in order to explore the range of positive shifts in the voltage dependence of activation that are possible. These Na_V_Ab/Na_V_1.7-VS2 chimeras were generated by grafting nucleotide sequences of S1-S2 and S3-S4 segments of Na_V_1.7 VS2 into homologous regions in Na_V_Ab. The cDNAs encoding Na_V_Ab/Na_V_1.7-VS2 chimeras were generated by overlap extension PCR (Heckman and Pease, 2007) for Na_V_Ab 7DII, and by gene synthesis for Na_V_Ab 7DII-4 and 7DII-5. These cDNAs were then amplified by PCR and cloned into the modified pIZT vector described previously (Wisedchaisri et al. 2019). Briefly, the pIZT vector (Life Technologies) was modified to include GFP followed by a self-cleaving P2A sequence under an OpIE2 promoter to facilitate identification of transfected Sf9 cells for electrophysiological recording, and Na_V_Ab and mutants were expressed as GFP-P2A-Na_V_Ab. Na_V_Ab 7DII-2 and 7DII-3 chimeras were derived from Na_V_Ab 7DII-4 by site-directed mutagenesis. Single, double, and triple residue substitutions in Na_V_Ab 7DII chimeras and Na_V_Ab were carried out by site-directed mutagenesis. All constructs were characterized by electrophysiology using transfected Sf9 cells as described previously (Wisedchaisri et al., 2019). For more experimental details on construction and functional characterization of these constructs, see McCord, E., Broadband tuning the voltage dependence of a sodium channel. PhD Dissertation, University of Washington*: https://digital.lib.washington.edu/researchworks/handle/1773/45249*.

### Electrophysiology

1.5-3 MΩ glass pipettes were used to patch-clamp fluorescent Sf9 cells, and sodium currents were recorded via whole-cell voltage clamp (EPC10, Pulse, HEKA), sampled at 250 kHz or 500 kHz, and filtered at 3, 8, or 10 kHz. Membrane capacitance ranged from 5-25 pF. Rs was below 10 MΩ, with a 2 μs lag and at least 70% compensation, or with voltage error below 10 mV. A P/4 capacitive transient subtraction protocol was used for ionic current, and P/5 was used for gating current measurements, taking care to measure the leak current well outside of the voltage range of gating charge movement. Standard recording solutions were as follows: Intracellular (in mM): NaCl (35), CsF (105), EGTA (10), HEPES (10); Extracellular (in mM): NaCl (140), MgCl_2_ (2), CaCl_2_ (2), HEPES (10). In order to analyze Na_V_Ab mutants with positively shifted voltage dependence, solutions were designed to result in reversal potentials outside the range of the voltage response by substituting NMDG-Cl for NaCl in both solutions as needed. For gating charge measurements, the intracellular solution contained NMDG-Cl (140), EGTA (10), HEPES (10), MgCl_2_ (1), and external contained NMDG-Cl (148), HEPES (10), MgCl_2_ (3). The pH for both intracellular and extracellular solutions was 7.4, adjusted with CsOH/HCl and NaOH/HCl for solutions that contained no NMDG-Cl, respectively, and NMDG+/HCl for those containing NMDG. The measured liquid junction potential was −7 mV for ionic current recordings, and the reported voltages were not adjusted. Ionic current families were generated by holding cells at −150 mV and stimulating with a series of 50-ms pulses at 10-mV intervals. All scale bars denote 1 nA/10 msec. To determine the reversal potential of outward current, peak current was elicited by pulsing to an appropriate potential for each construct and decreasing the voltage in 10 mV steps until substantial inward current was seen in the tail currents. An agar bridge consisting of extracellular solution with 3% Agar was placed between the reference electrode and the bath in experiments utilizing perfusion of extracellular solution containing reducing agents to isolate the electrode from the reducing agents.

### Data analysis

Data were analyzed using IGOR Pro (6.37). The peak current at each voltage of the current family was averaged and plotted as a function of the stimulus voltage to visualize the current vs voltage (I/V) relationship. When the I/V curves contained only outward current, the reversal potential (V_rev_) was determined from the tail currents at peak activation. The instantaneous current of the tail current family was plotted as a function of voltage, fit with a line near 0 mV, and the X intercept was used as V_rev_ to generate the conductance vs voltage (G/V) curve from the I/V curve by calculating G=I/(V_m_−V_rev_). Normalized G/V curves were fit with a simple two-state Boltzmann equation 1/(1+exp(V_1/2_−V_m_)/k) in which V_m_ is the stimulus potential, V_A_ is the half-activation voltage, and k is a slope factor. I/V plots of inward currents were fit with (V_m_−V_rev_)*(I_min_/(1+exp((V_m_−V_A_)/k)) to generate Boltzmann relationships fit to normalized G/V curves. Q/V curves were generated by integrating the P/N subtracted curve either during the test pulse or after the return to the prepulse voltage.

## RESULTS

We took a stepwise approach to determining whether broadband tuning of the voltage dependence of activation (V_A_) of a sodium channel is feasible. First, we asked whether formation of chimeric channels with amino acid residues from a mammalian sodium channel is able to induce positive shifts in the V_A_ of Na_V_Ab. Second, we surveyed the V_A_-shifting effects of single amino acid substitutions selected from the literature, which were known to induce positive shifts in V_A_ in other channel types. Third, we assessed the effects of pairwise and triple combinations of these mutations in chimeric mammalian Na_V_1.7/Na_V_Ab channels. Fourth, we determined if a selected triple-mutant combination was able to shift the V_A_ of Na_V_Ab itself and analyzed whether those positive shifts in V_A_ are accompanied by similar positive shifts in the voltage dependence of gating charge movement (V_Q_). To facilitate clear presentation of this large amount of data, we focus on measurements of V_A_ and V_Q_. Additional data on the kinetics of activation and inactivation and on steady-state inactivation of selected mutants are available in McCord, PhD Dissertation, University of Washington*: https://digital.lib.washington.edu/researchworks/handle/1773/45249*.

### Shifting Voltage Dependence of a Chimeric Sodium Channel

The bacterial sodium channel Na_V_Ab activates at very negative membrane potentials, with V_A_ = −98 mV (Gamal El-Din et al., 2013a). Mammalian Na_V_ channels activate at much more depolarized membrane potentials. For example, the Na_V_1.7 channel activates with a V_A_ of −22 mV (Wu et al., 2013). To test whether substitution of amino acid residues from mammalian Na_V_1.7 into the VS of Na_V_Ab could shift its voltage dependence, we constructed and analyzed chimeras of Na_V_Ab and Na_V_1.7 (Figure 1; Table 1). The basic chimera 7DII contained the extracellular residues of the human Na_V_1.7 DII VSD, starting near the extracellular end of each transmembrane helix (Figure 1A and C, left, Table 1). This construction shortened the S1-S2 liker by one residue and lengthened the S3-S4 linker by four residues, resulting in a net gain of three residues for the channel (Table 1). We measured the sodium current observed when this chimera was depolarized to a range of membrane potentials, and we calculated the conductance as a function of stimulus potential. The resulting conductance/voltage (G/V) curves for chimera 7DII revealed a similar voltage dependence of activation to Na_V_Ab (V_A_ = −118 ± 0.9 mV; Figure 1B, red; Table 2). In contrast to this small effect on voltage dependence, substitution of additional amino acid residues in chimera 7DII caused large positive voltage shifts. V_A_ shifted to −7.5 ± 1.7 mV for chimera 7DII-2 (Figure 1B, orange) and further shifted to +34.6 ± 1.4 mV for chimera 7DII-3 (Figure 1B, blue), a total voltage shift of +153 mV (Table 2). These large voltage shifts were caused by human Na_V_1.7 residues substituted for Na_V_Ab residues over the extracellular halves of the S1-S4 segments (Figure 1C). This dramatic positive shift of V_A_ suggested the possibility that the voltage dependence of chimera 7DII could be tuned over a broad voltage range by substitutions of individual amino acid residues and by pairwise and triple combinations of individual mutations.

**Figure 1.**
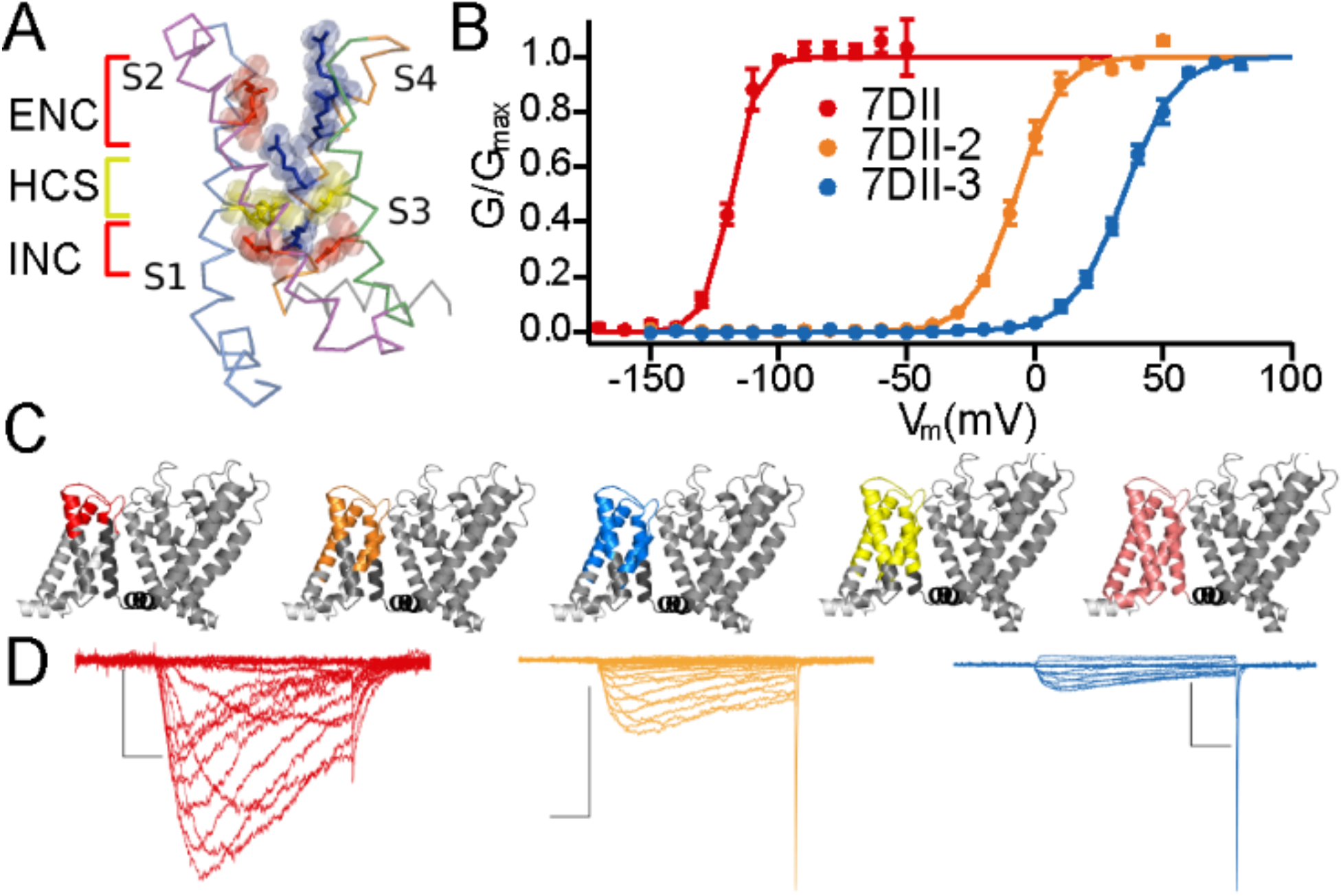
Voltage dependence of activation of Na_V_Ab and Na_V_Ab/Na_V_1.7 chimeras. **A.** Voltage sensor model highlighting key functional components: ENC, extracellular negative cluster (red); HCS, hydrophobic constriction site (gold); INC, intracellular negative cluster (red); gating charge Arg residues (blue). **B.** Conductance-voltage (G/V) relationships of three chimeras containing increasing numbers of human Na_V_1.7 residues. **C.** Homology models (Swiss-Model, 5X0M template (Shen et al., 2017), Domain II) were made to highlight the extent of substitution of human residues in five chimeric human-bacterial sodium channels created by replacing Na_V_Ab residues with those of human Na_V_1.7 DII. The S1-S4 segments are colored from light to dark gray, with the S4-S5 linker in black. The colored portion indicates human residues, while native Na_V_Ab residues are in varying shades of gray. The colors correspond to the different chimeric channels. The channels are arranged in order from least to most humanized: red, 7DII; orange, 7DII-2; blue, 7DII-3; yellow, 7DII-4; and pink, 7DII-5. The latter two constructs did not conduct sodium current. **D.** Current records of the three functional chimeras colored as indicated in B and C. Individual G/V curves were normalized to peak current values and fit with a standard Boltzmann equation. The average V_A_ and k values were used to draw the solid line fits. Datapoints for each construct represent average values at the indicated voltages, and error bars denote mean ± SEM.

**Table 1.**
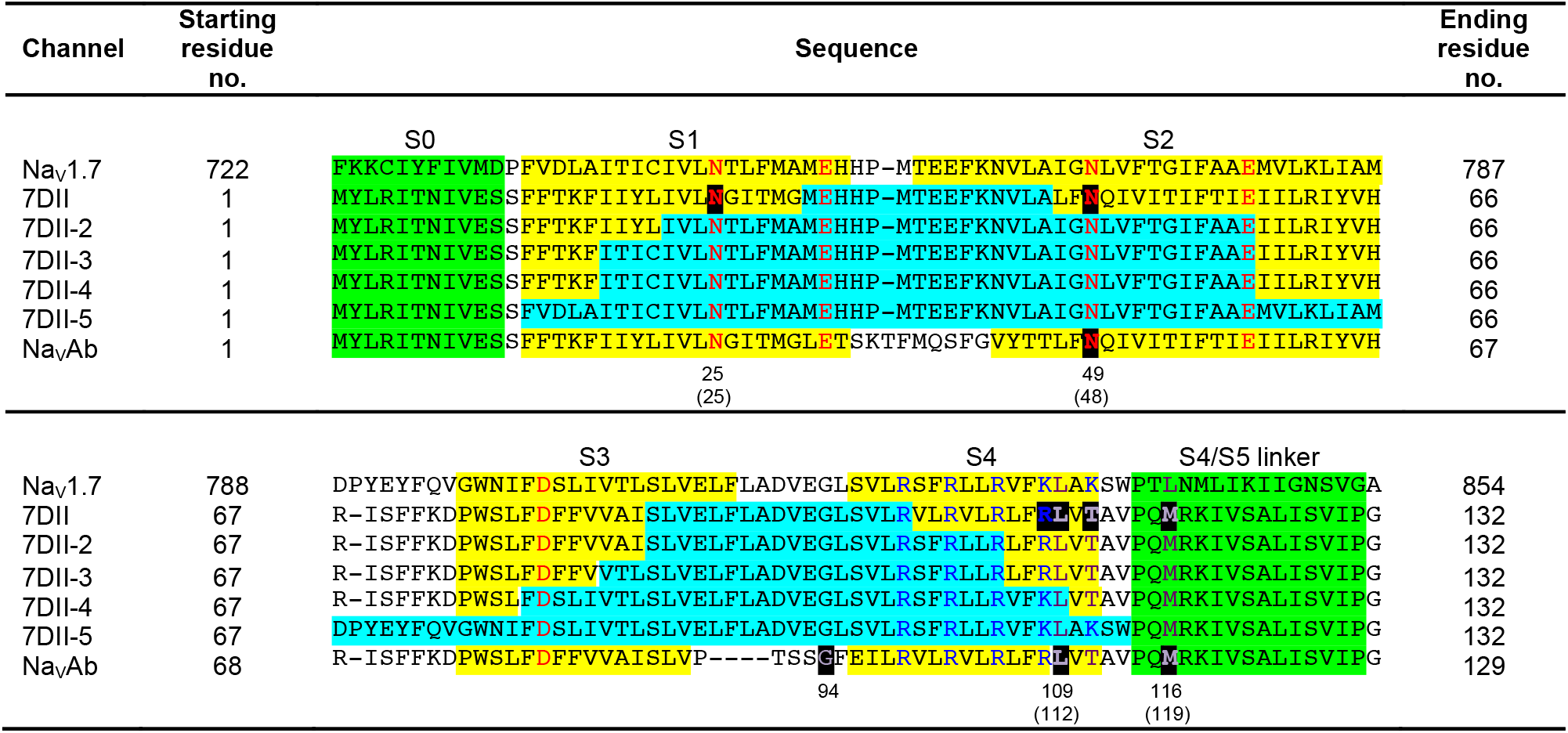
Sequence alignment of Na_V_1.7-DII, Na_V_Ab WT, and chimeras. Yellow highlighting indicates transmembrane segments. Green highlighting indicates intracellular helices. Cyan highlighting indicates human residues transplanted into Na_V_Ab to create the indicated chimeras. Intracellular and extracellular negative cluster residues in S1-S3 segments are shown in red. The arginine gating charges in S4 segment are shown in blue. As a landmark, mutations characterized in this study are highlighted in black background with residue numbers for Na_V_Ab and 7DII chimera (in parenthesis) shown beneath.

**Table 2.** Activation parameters of Na_V_1.7/Na_V_Ab-VS2 chimeras.

| Chimera | $V_A$ | k | $Z_{app}$ | n |
| --- | --- | --- | --- | --- |
| 7DII | $-118.0 \pm 0.9$ | $5.0 \pm 0.3$ | $5.1 \pm 0.4$ | 7 |
| 7DII-2 | $-7.5 \pm 1.7$ | $8.5 \pm 0.6$ | $2.9 \pm 0.2$ | 2 |
| 7DII-3 | $34.6 \pm 1.4$ | $10 \pm 0.5$ | $2.5 \pm 0.1$ | 6 |
| 7DII-4 | N.D. | N.D. | N.D. | N.D. |
| 7DII-5 | N.D. | N.D. | N.D. | N.D. |

$V_A$ , half maximal activation voltage; k, slope factor; $Z_{app}$ , apparent gating charge movement. Mean ± SEM. N.D., not determined.

### Shifting V_A_ with Single Mutations

Na_V_Ab has three distinct kinetic phases of slow inactivation: two that are easily reversible by hyperpolarization, and one “late use-dependent phase” that is induced by repetitive depolarizing pulses and reverses very slowly (Blanchet and Chahine, 2007; Gamal El-Din et al., 2013b; Pavlov et al., 2005). The late phase of inactivation makes stable recordings of voltage-dependent properties of Na_V_Ab channels impractical. Previous work showed that the mutation N49K (numbered N48K in the Na_V_Ab/Na_V_1.7 7DII chimera) caused a positive shift in V_A_ and blocked slow, use-dependent inactivation of Na_V_Ab (Gamal El-Din et al., 2013a). As expected from this prior work, the N48K mutation in chimera 7DII shifted 73 mV from V_A_ = −118 ± 0.9 mV to V_A_ = −45.4 ± 0.9 mV (Figures 2 and 3, red), and it eliminated the late use-dependent inactivation (not shown). This residue is located in a conserved site on the S2 segment containing an amino acid residue in the Extracellular Negative Cluster (ENC) that acts as a counter charge to the positive gating charges (Figure 2C). In Na_V_Ab, the native residue is Asn, which is hydrophilic and has a partial net negative charge. Placing a positive charge in this position may alter gating charge interactions and oppose both activation and slow, use-dependent inactivation.

**Figure 2.**
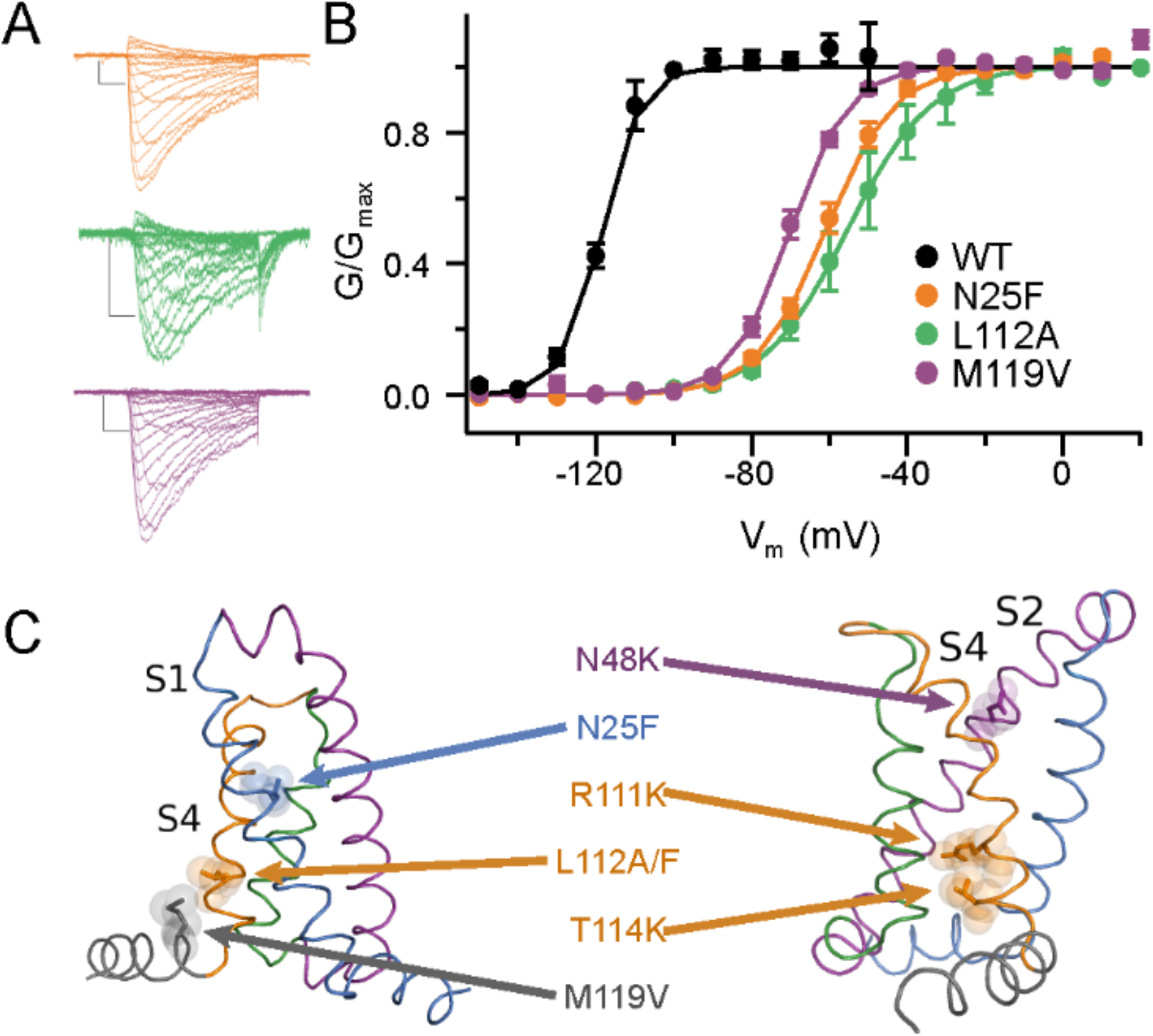
Functional effects of single uncharged residue substitutions. **A.** Representative current records. **B.** G/V relationships of Na_V_Ab 7DII with three separate hydrophobic substitutions in the VS: WT 7DII, black; M119V, purple; L112A, green; N25F, orange. **C.** Structural model of single residue substitutions. The color of the text corresponds to the coloring of the transmembrane segment in the model on which the residues are found: S1, blue; S2, purple; S3, green; S4, orange; S4-S5 Linker, gray. Individual G/V curves were normalized to peak current values and fit with a standard Boltzmann equation. The average V_A_ and k values were used to draw the solid line fits. Datapoints for each construct represent average values at the indicated voltage, error bars denote mean ± SEM.

**Figure 3.**
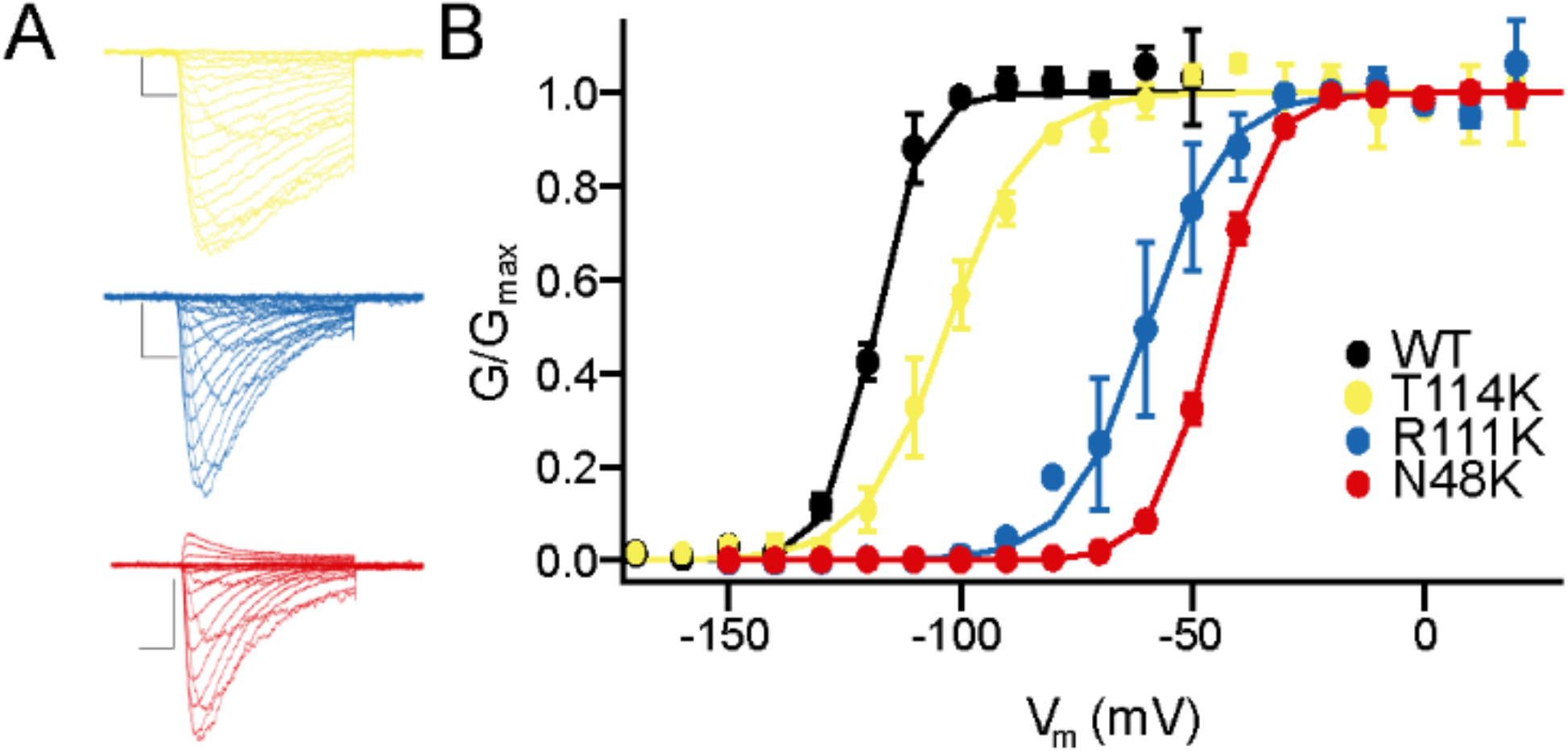
Functional effects of single charged residue substitutions. **A.** Representative current records. **B.** G/V relationships for 7DII alone and with three individual charged residue substitutions: WT 7DII black; T114K, yellow; R111K, blue; and N48K, red. Individual G/V curves were normalized to peak current values and fit with a standard Boltzmann equation. The average V_A_ and k values were used to draw the solid line fits. Datapoints for each construct represent average values at the indicated voltages, and error bars denote mean ± SEM.

To test whether other substitutions of single non-gating-charge residues would have comparable effects, we searched the literature for previously described, highly conserved residues whose mutation caused large positive shifts in V_A_. Mutagenesis studies of *Shaker B* potassium channels highlighted the residue equivalent to N25, located in the center of the VS, as a potential voltage-shifting amino acid substitution ((Lacroix et al., 2014); Figure 2C, Table 1). Mutation of N25 to Phe shifted the V_A_ by +57 mV to V_A_=−61.0 ± 1.4 mV (Figure 2A and B, orange; Table 3). Thus, this mutation has comparable effects on V_A_ in an insect potassium channel and a bacterial sodium channel.

**Table 3.** Activation parameters for 7DII single substitutions.

| Chimera Mutant | $V_A$ | k | $Z_{app}$ | n |
| --- | --- | --- | --- | --- |
| 7DII | $-118.0 \pm 0.9$ | $5.0 \pm 0.3$ | $5.1 \pm 0.4$ | 7 |
| N25F | $-61.0 \pm 1.4$ | $8.6 \pm 0.5$ | $2.9 \pm 0.2$ | 5 |
| N48K | $-45.4 \pm 0.9$ | $6.0 \pm 0.2$ | $4.2 \pm 0.2$ | 8 |
| N48K/Δ28 | $-47.8 \pm 1.8$ | $6.9 \pm 0.3$ | $3.6 \pm 0.2$ | 3 |
| R111K | $-59.8 \pm 6.6$ | $8.4 \pm 0.6$ | $3.0 \pm 0.02$ | 2 |
| L112A | $-55.0 \pm 4.9$ | $10.5 \pm 1.6$ | $2.4 \pm 0.4$ | 2 |
| T114K | $-102.0 \pm 2.9$ | $8.9 \pm 1.0$ | $2.9 \pm 0.4$ | 3 |
| M119V | $-70.0 \pm 1.2$ | $7.4 \pm 0.2$ | $3.3 \pm 0.1$ | 5 |

$V_A$ , half maximal activation voltage; k, slope factor; $Z_{app}$ , apparent gating charge movement. Mean ± SEM.

Continuing this line of investigation, we sought well-characterized mutations of conserved amino acid residues located near the intracellular end of S4 that cause positive shifts in V_A_. We first focused on Na_V_Ab/L109 (Table 1), whose counterpart in *Shaker B* potassium channels and Na_V_1.2 sodium channels yielded large positive shifts in V_A_ upon mutation (Kontis et al., 1997; Lopez et al., 1991). Mutation of the corresponding residue in 7DII (L112, Table 1, Figure 2C) to Ala caused a positive shift of V_A_ to −55.0 ± 4.9 mV in 7DII (Figure 2A and B, green; Table 3). Taking a similar approach, we found that mutation M119V of 7DII positively shifted V_A_ to −70.0 ± 1.2 mV (Figure 2A and B, purple; Table 3). L112A is located directly after R4, and M119 begins the S4-S5 linker (Figure 2C). Thus, these two amino acid residues flank the intracellular end of the S4 segment and may impede its movement during voltage sensing and gating.

These results suggested that the intracellular end of the S4 segment might be a hotspot for V_A_-shifting mutations. To follow up that idea, we analyzed the effects of a mutation of the R4 gating charge that preserves its positive charge (R111K) and a mutation of T114 (T114K; Tables 1 and 3) that introduces an additional positive charge in register with the R1-R4 gating charges. Substitution of these residues also resulted in positive shifts of V_A_, with R111K causing a +58-mV shift of V_A_ to −59.8 ± 6.6 mV (Figure 3, blue) and T114K causing a +16-mV shift of V_A_ to −102 ± 2.9 mV (Figure 3, yellow). Altogether, these experiments identify a set of conserved amino acid residues whose mutation individually causes substantial positive shifts in V_A_.

### Pairwise Combinations of V_A_ Shifting Mutations

Because these individual mutations gave promising results showing substantial positive shifts in V_A_, we analyzed the effects of pairwise combinations of selected V_A_-shifting mutations (Figure 4). The effects of the combined mutations R111K and T114K were greater than additive, as the total shift in V_A_ of the double mutant (+102 mV) was greater than the sum of the two individual shifts in V_A_ (+74 mV, Figure 4A and B, brown, Table 4). The combined shift of V_A_ of the N48K/R111K double mutant was +156 mV, which exceeds the sum of the individual shifts of +131 mV (Figure 4C, blue; Table 4). The combinatorial effects of mutation N48K with R111S or R111C on V_A_ were smaller than R111K, but also showed cumulative positive shifts in V_A_ (Figure 4C, yellow and green; Table 4). These results encouraged us to construct triple mutations to search for combinations that gave even larger positive shifts in V_A_.

**Figure 4.**
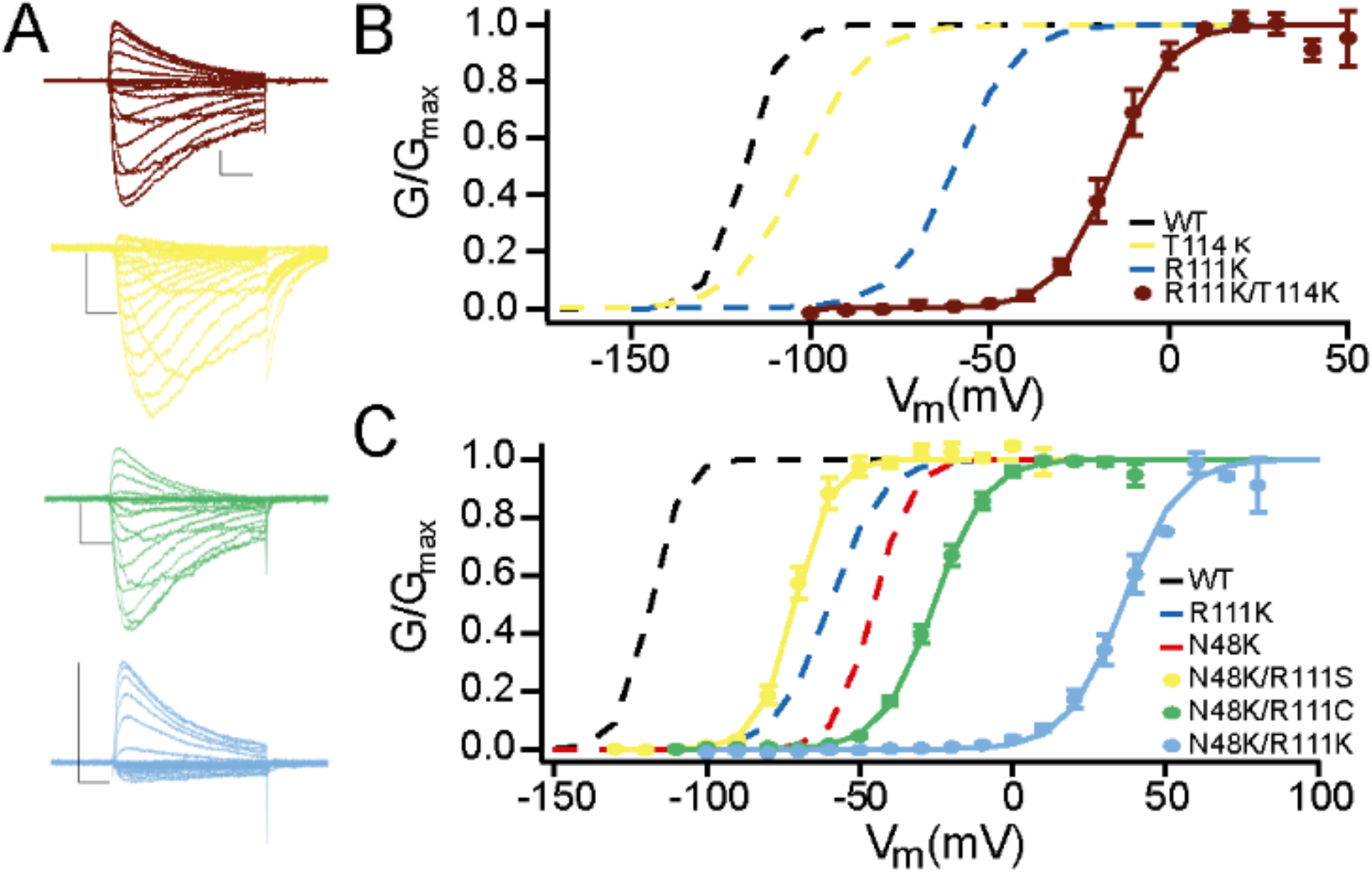
Combinatorial effects of paired mutations. **A.** Representative sodium currents. Current records are colored to correspond to their G/V curves in panel B. **B.** G/V relationships for WT 7DII, Na_V_Ab/R111K alone, blue; T114K alone, yellow; and R111K/T114K (brown). **C.** G/V relationships for WT 7DII and double mutants: N48K/R111S, yellow; N48K/R111C, green; N48K/R111K, light blue. Dashed traces for the indicated constructs are presented for comparison based on data in previous figures. Individual G/V curves were normalized to peak current values and fit with a standard Boltzmann equation. The average V_A_ and k values were used to draw the solid and dotted line fits. Datapoints for each construct represent average values at the indicated voltages, and error bars denote mean ± SEM. Dashed lines denote the average G/V fit of constructs described in previous figures.

**Table 4.**
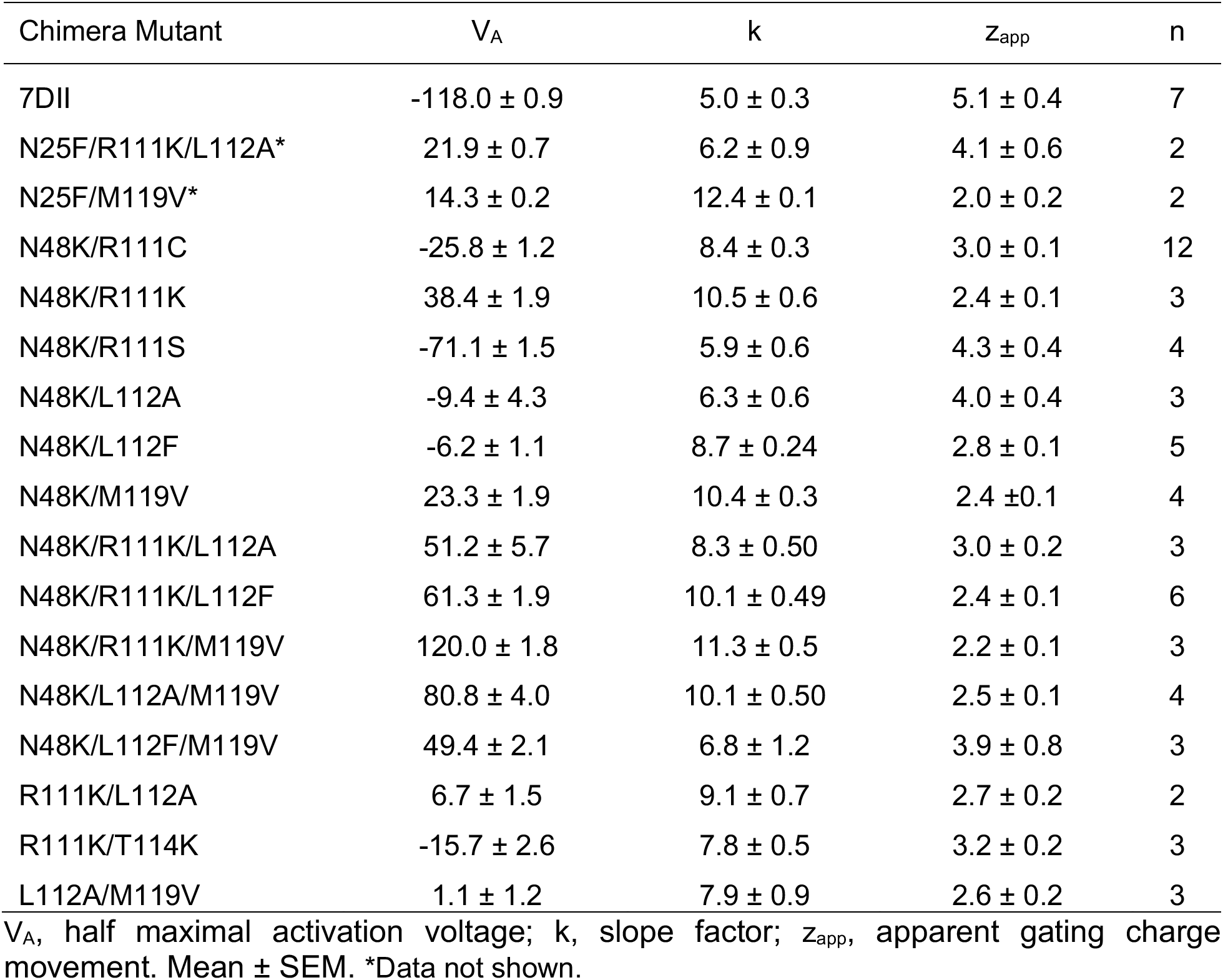
Activation parameters of 7DII and combinations of substitutions.

| Chimera Mutant | $V_A$ | k | $Z_{app}$ | n |
| --- | --- | --- | --- | --- |
| 7DII | $-118.0 \pm 0.9$ | $5.0 \pm 0.3$ | $5.1 \pm 0.4$ | 7 |
| N25F/R111K/L112A* | $21.9 \pm 0.7$ | $6.2 \pm 0.9$ | $4.1 \pm 0.6$ | 2 |
| N25F/M119V* | $14.3 \pm 0.2$ | $12.4 \pm 0.1$ | $2.0 \pm 0.2$ | 2 |
| N48K/R111C | $-25.8 \pm 1.2$ | $8.4 \pm 0.3$ | $3.0 \pm 0.1$ | 12 |
| N48K/R111K | $38.4 \pm 1.9$ | $10.5 \pm 0.6$ | $2.4 \pm 0.1$ | 3 |
| N48K/R111S | $-71.1 \pm 1.5$ | $5.9 \pm 0.6$ | $4.3 \pm 0.4$ | 4 |
| N48K/L112A | $-9.4 \pm 4.3$ | $6.3 \pm 0.6$ | $4.0 \pm 0.4$ | 3 |
| N48K/L112F | $-6.2 \pm 1.1$ | $8.7 \pm 0.24$ | $2.8 \pm 0.1$ | 5 |
| N48K/M119V | $23.3 \pm 1.9$ | $10.4 \pm 0.3$ | $2.4 \pm 0.1$ | 4 |
| N48K/R111K/L112A | $51.2 \pm 5.7$ | $8.3 \pm 0.50$ | $3.0 \pm 0.2$ | 3 |
| N48K/R111K/L112F | $61.3 \pm 1.9$ | $10.1 \pm 0.49$ | $2.4 \pm 0.1$ | 6 |
| N48K/R111K/M119V | $120.0 \pm 1.8$ | $11.3 \pm 0.5$ | $2.2 \pm 0.1$ | 3 |
| N48K/L112A/M119V | $80.8 \pm 4.0$ | $10.1 \pm 0.50$ | $2.5 \pm 0.1$ | 4 |
| N48K/L112F/M119V | $49.4 \pm 2.1$ | $6.8 \pm 1.2$ | $3.9 \pm 0.8$ | 3 |
| R111K/L112A | $6.7 \pm 1.5$ | $9.1 \pm 0.7$ | $2.7 \pm 0.2$ | 2 |
| R111K/T114K | $-15.7 \pm 2.6$ | $7.8 \pm 0.5$ | $3.2 \pm 0.2$ | 3 |
| L112A/M119V | $1.1 \pm 1.2$ | $7.9 \pm 0.9$ | $2.6 \pm 0.2$ | 3 |

$V_A$ , half maximal activation voltage; k, slope factor; $Z_{app}$ , apparent gating charge movement. Mean ± SEM. \*Data not shown.

### Triple Combinations of V_A_ Shifting Mutations

We studied substitutions of residue L112 with Ala and Phe in combination with N48K and R111K (Figure 5). When combined with N48K, L112A and L112F had similar V_A_ values of −9.4 ± 4.3 mV and −6.2 ± 1.1 mV, respectively, whereas the V_A_ for N48K alone was −45.4 ± 0.9 mV (Figure 5B, yellow and green; Table 3). With the further addition of R111K, the two triple mutants again had similar V_A_ values at +51.2 ± 5.7 mV for N48K/R111K/L112A and +61.3 ± 1.9 mV for N48K/R111K/L112F (Figure 5B, purple and orange). The shift between N48K and N48K/R111K was +84 mV (Figure 5A, Table 4). When R111K was added to N48K/L112A and N48K/L112F, the shifts were much smaller at +61 mV and +68 mV, respectively, suggesting that there is an opposing interaction between R111K and the two L112 substitutions (Figure 5B, purple and orange, Table 4). The triple mutant N48K/R111K/M119V produced the largest shift in V_A_ of all constructs tested, +240 mV, to give a V_A_ value of +120 ± 1.8 mV (Figure 5A, green, Table 4).

**Figure 5.**
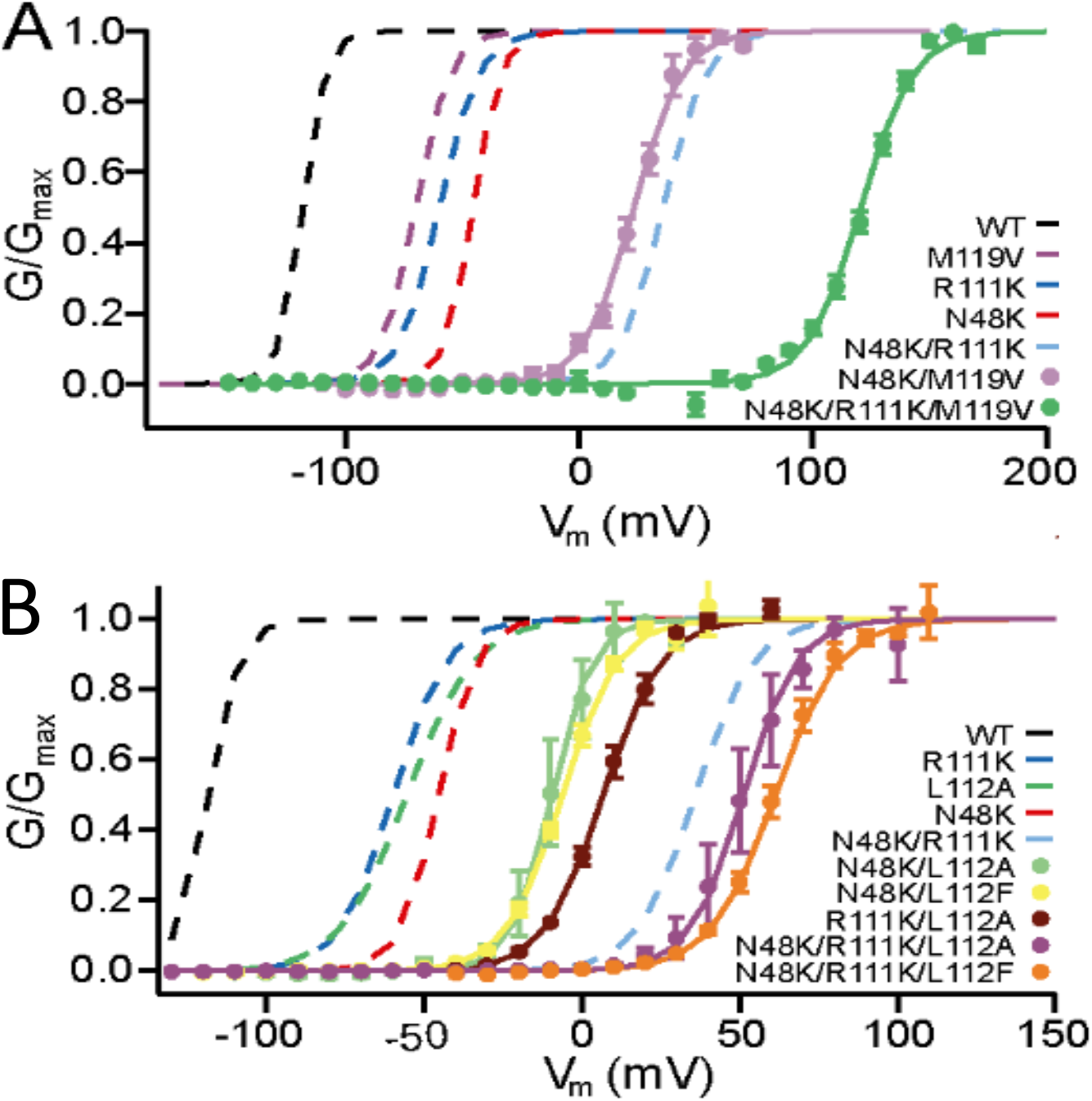
Combinations of N48K, R111K, and M119V or L112A/F. **A.** G/V relationships of WT 7DII (black), 7DII/N48K/M119V (lavender) and N48K/R111K/M119V (green). **B.** G/V relationships of WT 7DII (black), 7DII/N48K/L112A (green), N48K/L112F (yellow), R111K/L112A (brown), N48K/R111K/L112A (purple), and N48K/R111K/L112F (orange). Dashed traces for the indicated constructs are presented for comparison based on data in previous figures. Individual G/V curves were normalized to peak current values and fit with a standard Boltzmann equation. The average V_A_ and k values were used to draw the solid and dotted line fits. Datapoints for each construct represent average values at the indicated voltages, and error bars denote mean ± SEM. Dashed lines denote the average G/V fit of constructs described in previous figures.

We further explored triple combination mutations that did not include the gating charge R4 (R111). The double mutation N48K/L112A shifted V_A_ to −9.4 ± 4.3 mV, whereas the double mutation N48K/M119V shifted V_A_ to +23.3 ± 1.9 mV (Figure 6, light green and purple). Remarkably, the triple mutation N48K/L112F/M119V shifted V_A_ by +170 mV (Figure 6, brown, Table 4) and the triple mutation N48K/L112A/M119V shifted V_A_ by +200 mV, all the way to V_A_ = +80.8 ± 4.0 mV, without mutation of the R4 gating charge (Figure 6, orange, Table 4). These results demonstrate that the electrostatic force exerted on the gating charges by the transmembrane electrical field can be overcome by only three amino acid substitutions, which do not change the charge of the affected amino acid residues yet substantially alter the energetic landscape due to the chemical energy gained from new protein interactions.

**Figure 6.**
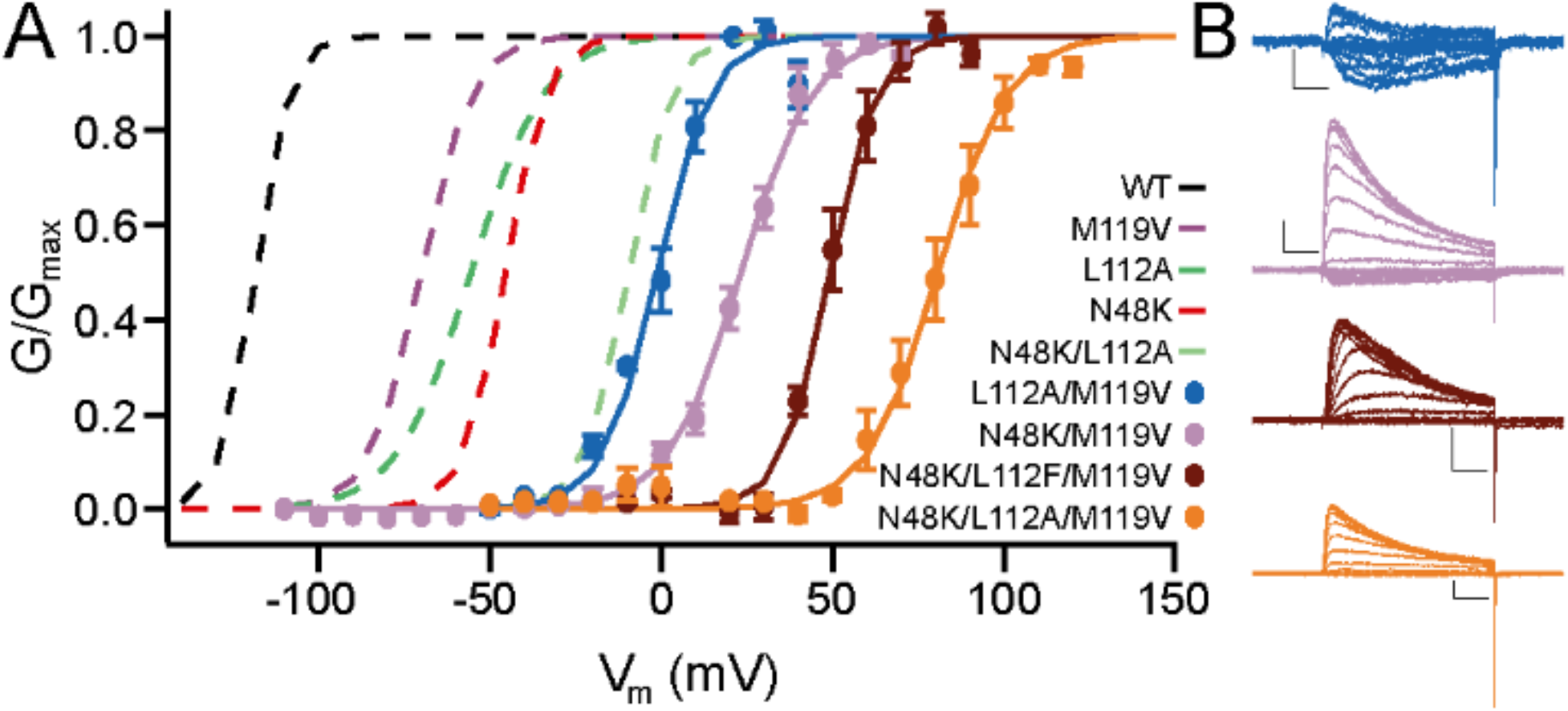
Voltage dependence of activation of 7DII with combinations of N48K, L112A, and M119V (KAV). **A.** G/V relationships for the 7DII chimeras with single, double, and triple combinations of N48K, M119V, L112A. WT 7DII (black dashes); M119V (purple dashes); L112A (green dashes); N48K (red dashes); N48K/L112A (light green dashes); L112A/M119V (blue); N48K/M119V (purple); N48K/L112F/M119V (brown); N48K/L112A/M119V (orange). Individual G/V curves were normalized to peak current values and fit with a standard Boltzmann equation. The average V_A_ and k values were used to draw the solid and dotted line fits. Datapoints for each construct represent average values at the indicated voltages, and error bars denote mean ± SEM. Dashed lines denote the average G/V fit of constructs described in previous figures. **B.** Representative current records are colored to correspond to their respective G/V curves. From top to bottom they are 7DII/L112A/M119V, N48K/M119V, N48K/L112F/M119V, and N48K/L112A/M119V.

### Positive Shifts in the Voltage Dependence of Gating Charge Movement

Our extensive studies of broadband tuning of the voltage dependence of activation of the bacterial/human chimera 7DII showed that its voltage sensitivity could be shifted cumulatively by up to +240 mV (Figure 5A, green). However, V_A_ is an indirect measure of movement of the S4 gating charges. In order to compare the effects of our V_A_-shifting mutations on both V_A_ and the voltage dependence of gating charge movement, V_Q_, we used the Na_V_AbΔ28 construct that we developed for high-level expression for structural studies (Lenaeus et al., 2017), in which 28 nonconserved amino acid residues are deleted from the C-terminus. This construct allowed introduction of combined V_A_-shifting mutations with high expression. We focused on combination mutations that did not include the R4 gating charge. Introduction of the combination of N48K, L112A, and M119V equivalent mutations (KAV) into Na_V_Ab caused a substantial positive shift in the V_A_ of Na_V_Ab (Figure 7A, green; Table 5). The N48K equivalent mutation alone caused a +75 mV shift of V_A_ in Na_V_Ab (Figure 7A, blue). The combined KAV mutation further shifted activation positively, to a V_A_ of +61.3 ± 2.1 mV (Figure 7A, green). Importantly, there was no detectable sodium current activated at 0 mV (Figure 7A, green). Thus, the Na_V_AbΔ28/KAV channel functions entirely in the positive voltage domain.

**Figure 7.**
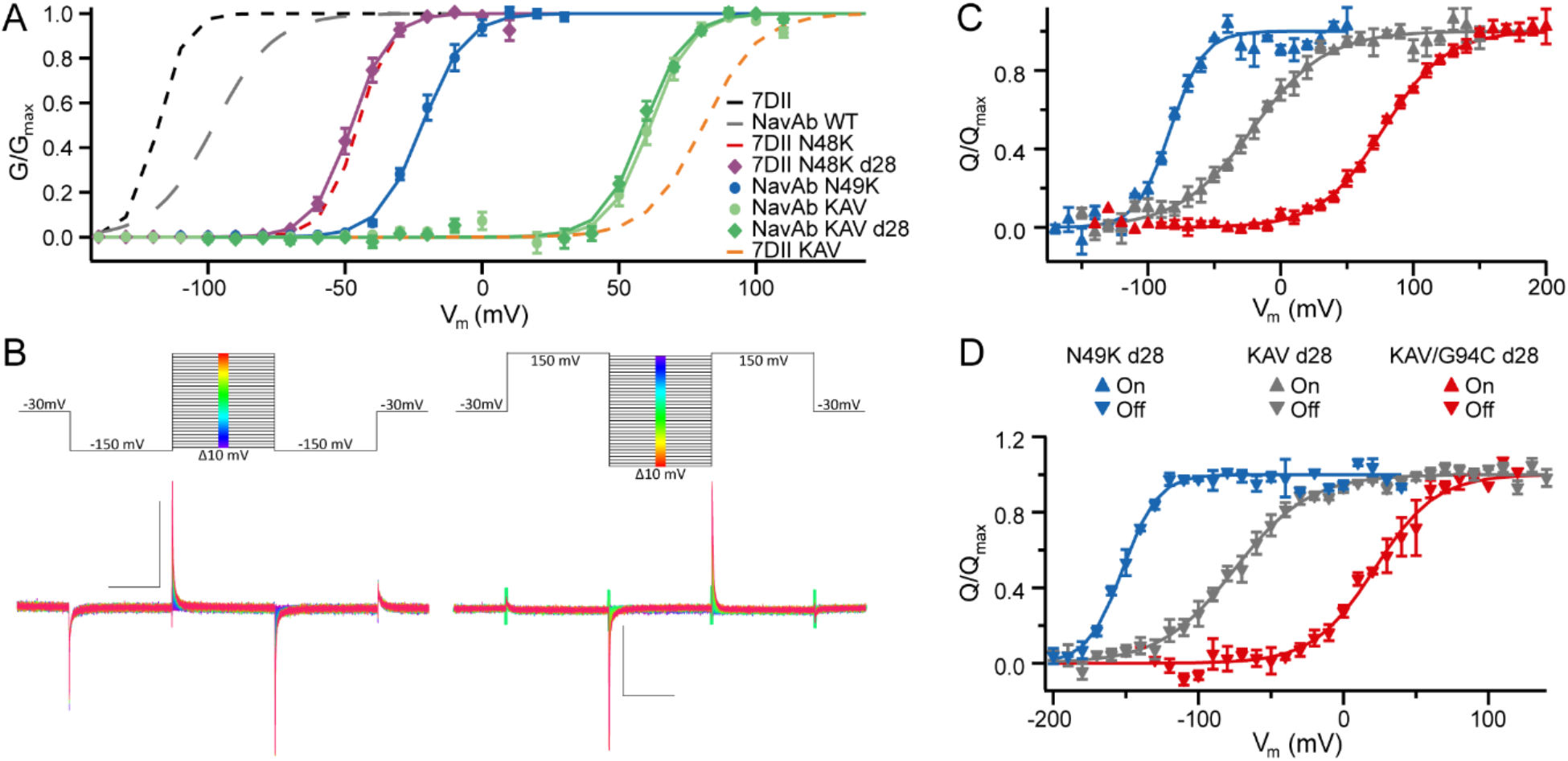
Comparison of the effects of the KAV mutations on G/V and Q/V relationships. **A.** G/V relationships for the indicated constructs: 7DII (black dashes); Na_V_Ab (gray dashes); 7DII/N48K (red dashes); 7DIIΔ28/N48K (purple diamonds); Na_V_Ab/N49K (blue circles); Na_V_Ab N49K/L112A/M1119V (KAV, light green circles); Na_V_AbΔ28/KAV (green diamonds), and 7DII/KAV (orange dashes). **B.** Voltage command diagrams and representative gating current traces of Na_V_AbΔ28/KAV for ON (left) and OFF (right) gating current measurements. Calibration bars indicate 1 nA and 10 msec. **C.** ON gating charge measurements in response to depolarization in media that contained NMDG-Cl substituted for NaCl and CaCl_2_. Cells were briefly hyperpolarized, and the subsequent current step was increased in 10 mV increments. **D.** OFF gating charge movement in response to hyperpolarization following brief depolarizating prepulses. A P/5 leak subtraction was used at voltages that remained well outside the range of charge movement in either the positive or negative direction. The area under the curve was calculated and plotted as a function of voltage. When necessary, traces were baseline corrected to the current remaining at the end of the test pulse. Individual Q/V curves were fit with a standard Boltzmann equation. The average V_Q_ and k values were used to draw the solid line fits. Datapoints for each construct represent average values at the indicated voltage. Error bars denote mean ± SEM. KAV d28 denotes Na_V_Ab/KAV/Δ28.

**Table 5.** Activation parameters of Na_V_Ab/N49K and KAV constructs.

| $Na_vAb$ Mutant | $V_A$ | k | $Z_{app}$ | n |
| --- | --- | --- | --- | --- |
| N49K | $-21.8 \pm 1.8$ | $7.9 \pm 1.0$ | $3.2 \pm 0.3$ | 4 |
| KAV | $61.3 \pm 2.1$ | $7.2 \pm 0.4$ | $3.4 \pm 0.2$ | 3 |
| KAV/Δ28 | $59.2 \pm 0.8$ | $7.7 \pm 0.8$ | $3.3 \pm 0.3$ | 3 |
| KAV/G94C/Δ28 | $88.4 \pm 1.1$ | $10.5 \pm 0.4$ | $2.3 \pm 0.1$ | 3 |
| KAV/Q150C | $77.8 \pm 0.8$ | $9.8 \pm 0.3$ | $2.5 \pm 0.7$ | 3 |
| KAV/Q150C/Δ28 | $76.7 \pm 3.1$ | $10.0 \pm 0.9$ | $2.5 \pm 0.2$ | 4 |
| KAV/G94C/Q150C/Δ28 | $105.0 \pm 0.8$ | $10.2 \pm 0.2$ | $2.4 \pm 0.6$ | 3 |

$V_A$ , half maximal activation voltage; k, slope factor; $Z_{app}$ , apparent gating charge movement. Mean ± SEM.

Na_V_Ab is Cys-free, and its relative, NaChBac, has been extensively studied by insertion of Cys residues and subsequent crosslinking by disulfide bond formation (Yarov-Yarovoy et al., 2012). We reasoned that introduction of carefully selected Cys residues might further shift V_A_ and V_Q_. As described previously (Wisedchaisri et al., 2019), we identified constructs with pairs of substituted Cys residues that expressed well, retained functional activity, and formed disulfide bonds in the resting state in a voltage-dependent manner. Surprisingly, we found even more positive shifts in V_A_ when single Cys residues were substituted for G94 and Q150 (Table 5; (Wisedchaisri et al., 2019)). G94C and Q150C shifted the voltage dependence of activation of Na_V_Ab/KAV positively to V_A_ = +88.4 ± 1.1 mV and +76.7 ± 3.1 mV, respectively. The double mutant, analyzed under in reducing conditions to prevent disulfide bond formation, exhibited a V_A_ of +105 ± 0.8 mV (Table 5; (Wisedchaisri et al., 2019)). Importantly, these channels do not open at 0 mV, but the question of whether their voltage sensors remain in their functionally resting position at that voltage remained to be addressed.

The outward movement of the gating charges can be measured as a capacitative current as shown in classical voltage-clamp studies of the squid giant axon (Armstrong and Bezanilla, 1973). By measuring the outward (ON) movement of the S4 gating charges in response to incremental voltage increases from a very negative membrane potential (Figure 7B), we found that V_Q_ was strongly shifted to positive membrane potentials by these V_A_-shifting mutations (Figure 7C). The V_Q_ values for these constructs were −83.1 ± 0.7 mV for Na_V_Ab/N48K, −21.5 ± 3.7 mV for Na_V_Ab/KAV, and +77.2 ± 0.4 mV for Na_V_Ab/KAV/G94C. These results represent a shift of +62 mV with the addition of the ‘AV’ mutations to N49K, and +99 mV for the subsequent addition of G94C, for a total Q/V shift of +160 mV (Figure 7C, Table 6). For comparison, the same substitutions shifted the G/V curve by only +127 mV in total (Table 6). Thus, these results show that both gating charge movement and channel activation are prevented at 0 mV in these mutant constructs. Evidently, the positive shifts in V_A_ values are driven primarily by positive shifts in V_Q_ values.

**Table 6.** On and Off gating charge parameters of Na_V_Ab/N49K and KAV mutants.

| NavAb Mutant | On $V_Q$ | k | $Z_{app}$ | n |
| --- | --- | --- | --- | --- |
| N49K/Δ28 | $-83.1 \pm 0.7$ | $12.5 \pm 0.1$ | $2.0 \pm 0.01$ | 2 |
| KAV/Δ28 | $-21.5 \pm 3.7$ | $27.8 \pm 2.5$ | $0.91 \pm 0.85$ | 4 |
| KAV/G94C | $77.2 \pm 0.4$ | $23.69 \pm 2.3$ | $1.07 \pm 0.11$ | 4 |
| | Off $V_Q$ | k | $Z_{app}$ | n |
| N49K/Δ28 | $-152.0 \pm 0.6$ | $12.1 \pm 0.5$ | $2.0 \pm 0.1$ | 2 |
| KAV/Δ28 | $-74.7 \pm 6.0$ | $25.6 \pm 2.1$ | $1.0 \pm 0.1$ | 4 |
| KAV/G94C/Δ28 | $21.4 \pm 3.3$ | $21.0 \pm 3.8$ | $1.5 \pm 0.3$ | 3 |
| KAV/Q150C/Δ28 | $-54.0 \pm 4.7$ | $37.6 \pm 4.0$ | $0.7 \pm 0.1$ | 2 |

$V_Q$ , half maximal activation voltage of gating charge movement; k, slope factor; $Z_{app}$ , apparent gating charge movement. Mean ± SEM.

The OFF gating charge movement occurred at more negative potentials for all constructs, as expected, because of charge immobilization during inactivation that must be reversed by hyperpolarization. The V_Q_ for OFF gating charge movement for Na_V_Ab/N49K was −152 ± 0.6 mV, and the addition of ‘AV’ in the KAV mutant increased this by +77 mV to a V_Q_ of −74.7 ± 6.0 mV (Figure 7D, Table 6). Addition of G94C resulted in a further +96 mV shift of V_Q_ to +21.4 ± 3.3 mV, representing a total shift of +173 mV (Figure 7D, Table 6). No gating charge movement was observed for KAV/G94C/Q150C in the absence of dithiothreitol to reduce disulfide bonds (n=4, data not shown), which supports our conclusion that disulfide-locking of G94C to Q150C prevents S4 translocation.

## DISCUSSION

Our aim in this project was to explore whether broadband tuning of the voltage dependence of a voltage-gated sodium channel is possible. Sodium channels are in the resting state at negative membrane potentials and are activated by depolarization of the membrane to more positive membrane potentials. Although previous work had identified mutations that shift V_A_ significantly, those experiments had not tested the extent of voltage shifting that is possible by combinations of mutations. Here we combined previously described mutations in multiple voltage-gated ion channels in order to test the range of positive shifting of V_A_ that is possible. We were surprised to find that combination of a small number of mutations was able to shift V_A_ up to 240 mV and yet retain normal sodium channel expression and function. Our findings show that broadband shifting of V_A_ of a sodium channel is possible and provide a toolbox of methods and constructs that enable exploration of sodium channel states, functions, and structure in the positive membrane potential domain that was previously inaccessible to detailed biophysical and structural studies. The utility of this approach has been proven by use of one of these constructs to lock Na_V_Ab in its resting state and determine its structure at high resolution (Wisedchaisri et al., 2019). In addition, our results lead to several important insights into the voltage sensing and voltage-dependent gating of sodium channels as discussed below.

### Voltage dependence of gating of mammalian Na_V_ channels can be transferred to Na_V_Ab

Sodium channels in bacteria have V_A_’s in the range of −130 mV, consistent with the negative membrane potential of bacteria in the range of −160 mV. In contrast, mammalian sodium channels have V_A_’s in the range of −50 mV to −20 mV, consistent with the resting membrane potentials of nerve and muscle cells of −90 mV to −70 mV. Despite this large difference in voltage sensitivity, the VS of bacterial and mammalian sodium channels are very similar in structure, with RMSD values of <3 Å in their core transmembrane domains (eg., (Jiang et al., 2020)). Our results show that changes of a small number of amino acid sidechains by mutation is sufficient to give this large difference on voltage-dependent function. Thus, in evolution, it was possible to tune VS function over a broad range without significant structural change.

### Single V_A_ shifting mutations from other ion channels can be functionally transferred into Na_V_Ab

The similarity of structure of bacterial and mammalian sodium channel VS suggests that mutations that alter conserved amino acid residues across the broad range of ion channels might have similar effects on voltage dependence. Our results show that mutations initially described in *Drosophila* potassium channels, mammalian sodium channels, and other distantly related ion channels have similar effects when transferred into the bacterial sodium channel Na_V_Ab by site-directed mutations. Evidently, the structural basis for VS function is conserved across phylogeny, and only a small number of molecular changes are required to elicit dramatic changes in voltage dependence.

### Combined mutations have additive effects on voltage-dependent activation

Although individual mutations have been frequently shown to shift the V_A_ of sodium channels, an exhaustive study of additive effects of combined mutations that shift V_A_ has not previously been carried out. Many combination mutations we studied showed increases in V_A_ that were greater than either individual mutation alone, combined increases that were approximately additive, or increases in V_A_ that were even more than additive when considered simply as the sum of V_A_. Thus, a majority of these mutations worked together to enhance positive shifts in V_A_ rather than to oppose those shifts, as judged by this empirical procedure.

### Combined mutations can shift sodium channel function entirely into the positive membrane potential domain

We found that multiple triple combinations of mutations are able to shift V_A_ entirely into the positive membrane potential domain. For these triple mutations, essentially all individual sodium channels are in the resting state at 0 mV. These results show that a negative transmembrane potential exerting an electrostatic force on the positive gating charges to pull them inward is not necessary for formation of the resting state at 0 mV. Moreover, application of stimuli in the positive voltage range lead to voltage sensor activation and pore opening. Evidently, the chemical energy of altered protein interactions caused by our mutations is sufficient to pull the gating charges into their resting conformation without a negative resting membrane potential, from which they can be activated by imposing positive voltage stimuli.

### V_A_-shifting mutations also shift the voltage dependence of gating charge movement

In principle, mutations might shift V_A_ by impeding the outward movement of the gating charges in response to depolarization, preventing the coupling of gating charge movement to pore opening, or a combination of both effects. Importantly, for the mutations we studied most completely, V_Q_ measured from gating currents was shifted to a comparable or even greater extent than V_A_. These studies support the conclusion that the gating charges are locked in place in their resting state by the chemical energy of protein interactions contributed by the mutated amino acid residues in our constructs until the membrane potential is depolarized to 0 mV or beyond.

### Stabilizing sodium channel functional states

All structures of native wild-type bacterial and eukaryotic sodium channels have activated or partially activated voltage sensors because there is no membrane potential in a protein crystal or on a cryo-EM grid. Determination of the structures of other sodium channel states will require stabilizing them at 0 mV. Our results provide a toolbox of methods and mutant cDNAs that can stabilize specific functional states at 0 mV for structural studies. We have already used the KAV triple mutant and disulfide locking of substituted Cys residues to stabilize the resting state of Na_V_Ab and determine its structure by cryo-EM (Wisedchaisri et al., 2019). The structure of the resting state revealed remarkable conformation changes, including inward movement of the S4 segment and gating charges by 11.5 Å, which pushes the S4-S5 linker inward into the cytosol to form a sharply bent elbow. These conformational changes poise the resting state for rapid activation upon depolarization. Other mutations described here can in principle stabilize intermediate states of the voltage sensor to capture multiple resting states, partially activated intermediate states, and inactivated states. These approaches offer a pathway toward understanding the complete conformational cycle of voltage-gated sodium channel function.

## ACKNOWLEDGEMENTS

We thank Dr. Tamer Gamal El-Din (Pharmacology, University of Washington) for valuable discussion and comments on the manuscript. This research was supported by National Institutes of Health Research Grants R01 NS15751 and R35 NS111573 to W.A.C. and Research Training Grant T32 GM007750 to E.M.

